# PIRATE: A fast and scalable pangenomics toolbox for clustering diverged orthologues in bacteria

**DOI:** 10.1101/598391

**Authors:** Sion C. Bayliss, Harry A. Thorpe, Nicola M. Coyle, Samuel K. Sheppard, Edward J. Feil

**Affiliations:** The Milner Centre for Evolution, Department of Biology and Biochemistry, University of Bath, Bath BA2 7AY

**Keywords:** Microbial genomics, pangenomics, next-generation sequencing, bioinformatics

## Abstract

Cataloguing the distribution of genes within natural bacterial populations is essential for understanding evolutionary processes and the genetic basis of adaptation. Here we present a pangenomics toolbox, PIRATE (Pangenome Iterative Refinement And Threshold Evaluation), which identifies and classifies orthologous gene families in bacterial pangenomes over a wide range of sequence similarity thresholds. PIRATE builds upon recent scalable software developments to allow for the rapid interrogation of thousands of isolates. PIRATE clusters genes (or other annotated features) over a wide range of amino-acid or nucleotide identity thresholds and uses the clustering information to rapidly classify paralogous gene families into either putative fission/fusion events or gene duplications. Furthermore, PIRATE orders the pangenome using a directed graph, provides a measure of allelic variation and estimates sequence divergence for each gene family. We demonstrate that PIRATE scales linearly with both number of samples and computation resources, allowing for analysis of large genomic datasets, and compares favorably to other popular tools. PIRATE provides a robust framework for analysing bacterial pangenomes, from largely clonal to panmictic species.

**Availability:** PIRATE is implemented in Perl and is freely available under an GNU GPL 3 open source license from https://github.com/SionBayliss/PIRATE.

**Supplementary Information:** Supplementary data is available online.

## Background

For most bacteria the complement of genes for a given species is far greater than the number of genes in any one strain. Comprising core genes shared by all individuals in a species and accessory genes that are variously present or absent, the pangenome represents a pool of genetic variation that underlies the enormous phenotypic variation observed in many bacterial species. Through horizontal gene transfer, bacteria can acquire genes from this pangenome pool that bestow important traits such as virulence or antimicrobial resistance [1].

Over the last decade, advances in whole genome sequencing technologies and bioinformatic analyses have allowed the cataloguing of genes and intergenic regions that make up the pangenomes of many species [2–8].

Current approaches define genes on the basis of strict sequence identity thresholds [2,3,7,8], e-value cutoffs [5,6] and bit score ratios [4]. However, genes accrue variation at different rates under the influence of positive and purifying selection [9]. Therefore, it is difficult to define a single identity threshold beyond which genes cease to belong to the same family. Relaxed thresholds risk over-clustering of related gene families, whilst conservative thresholds risk over-splitting, by misclassifying highly divergent alleles of the same gene into multiple clusters. Over-splitting is likely to be especially problematic in vertically acquired core genes that have undergone strong diversifying selection or horizontally acquired accessory genes from multiple source populations which share a distant common ancestor. This can lead to misleading impressions of pangenome size and composition.

In order to evaluate and classify genetic diversity within the pangenome we have created the Pangenome Iterative Refinement And Threshold Evaluation (PIRATE) toolbox. PIRATE provides the means to create pangenomes from any annotated feature (e.g. CDS, tRNA, rRNA) over a user-defined range of amino acid or nucleotide identity thresholds. PIRATE provides measures of sequence divergence and allelic diversity within the sample. PIRATE also categorises paralogs into duplication and/or fission loci, loci disrupted by an insertion, deletion or nonsense mutation, providing additional context on gene provenance. This rapid, scalable method allows for a comprehensive overview of gene content and allelic diversity within the pangenome.

## Methods

### Pangenome Construction

The PIRATE pipeline has been summarised as a schematic in Figure 1.A. The input is a set of GFF3 files. Feature sequences are filtered and the dataset is reduced by iterative clustering using CD-HIT [2,10]. The longest sequence from each CD-HIT cluster is used as a representative for sequence similarity searching (BLAST/DIAMOND) [11,12]. The normalised bit scores of the resulting all-vs-all comparisons are clustered using MCL after removing hits which fall below a relaxed threshold of percentage identity (default: 50%) [13]. The initial clustering at this lower bounds threshold is used to define putative ‘gene families’ (Figure 1.B). Initial designations may not represent the final outputs as families containing paralogs maybe subsequently split during the paralog splitting step. MCL clustering is repeated over a range of user specified percentage identity thresholds (default 50-95% amino acid identity, increments of 5). Unique MCL clusters at higher thresholds are used to identify ‘unique alleles’ (Figure 1.B). Loci may be shared between multiple unique alleles (MCL clusters) at different percentage identity thresholds (e.g. Figure 1.B – Family B). PIRATE uses the highest threshold at which a ‘unique allele’ is observed to define the shared percentage identity in the resulting outputs.

**Figure 1.**
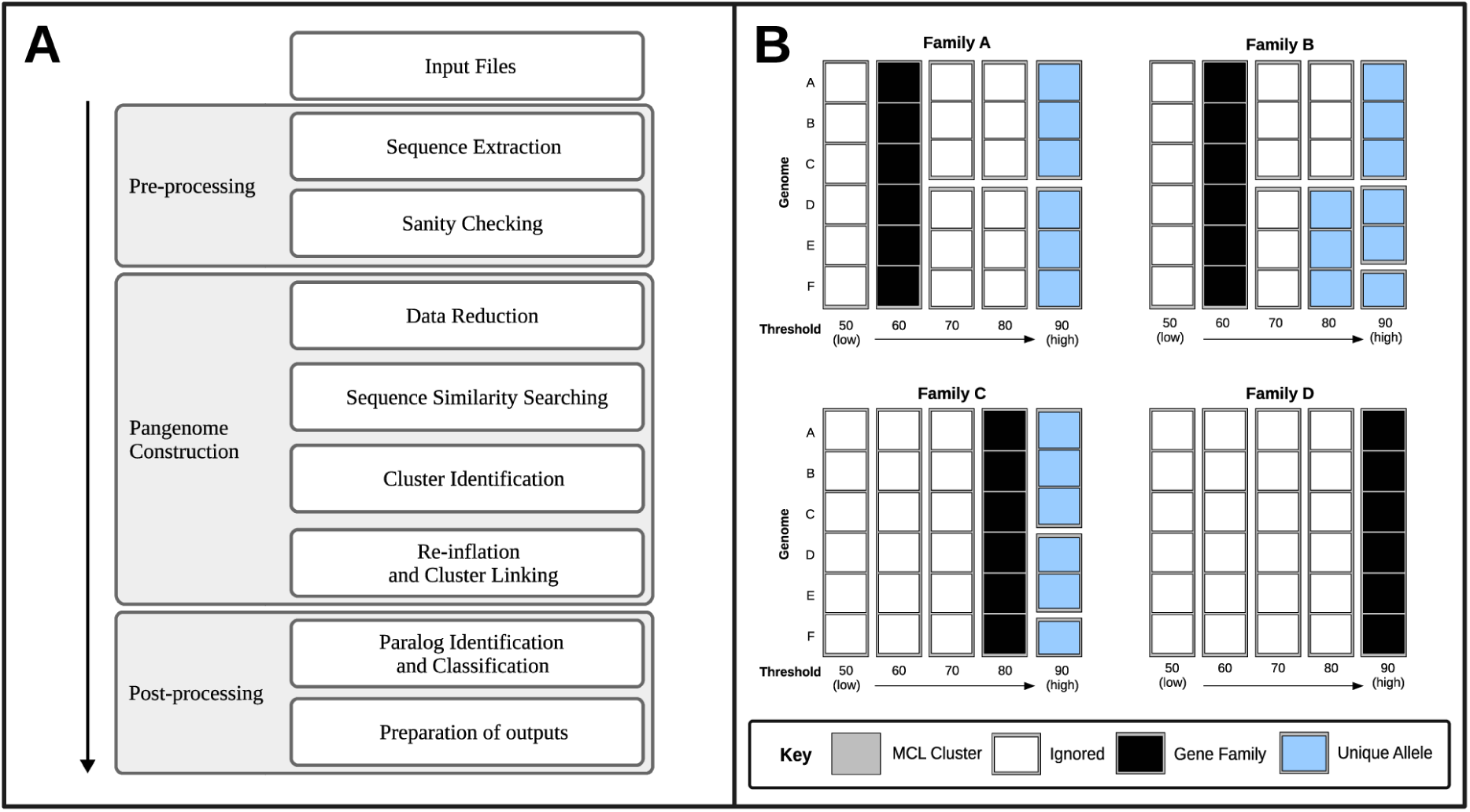
(A) Flow chart denoting a simplified workflow. (B) Example cluster classification. Blocks represent sequences from unique genomes. Grey blocks represent MCL clusters at various percentage identity cut-offs. Black squares indicate a ‘gene family’ cluster, the lowest %id threshold from the MCL clustering. Blue squares represent ‘unique alleles’, MCL clusters at higher % identity thresholds with unique combinations of sequences (at the higher threshold at which they are observed together). White squares represent redundant MCL clusters, these are not present in the PIRATE output.

### Paralog Classification

Clusters which contain more than one sequence per individual genome are putative paralogs and undergo an additional post-processing step (Supplementary Figure 6). All loci are clustered on the basis of sequence length (98% similar) using CD-HIT. Homology between representative loci is established using all-vs-all BLAST. Loci with no significant overlaps are considered putative fission loci and are compared against a reference sequence (the longest sequence in the gene family) which is considered the most ‘complete’ version of the gene. All combinations of putative fission loci are compared to the reference in order to find the combination which gives the most parsimonious coverage of the reference sequence. This combination locus is classified as a ‘fission locus’ that may have formed via gene disruption (e.g. insertion, deletion or nonsense mutation). Any locus which overlaps with all other loci or is not a part of a fission cluster is considered a duplication. The process is iterated until all loci have been classified.

### Cluster Splitting

After paralog classification, fission loci are treated as a single locus. Gene families that contain genomes with multiple loci, after accounting for fission loci, potentially represent two or more related gene families that have been over-clustered. In these cases the gene family is checked against the presence of MCL clusters (unique alleles) which contains a single copy of the loci in all constituent genomes (Supplementary Figure 6). These alleles are thereafter considered separate gene families with nomenclature denoting their shared provenance (e.g. g0001_1, g0001_2).

### Post-processing

Syntenic connections between gene families in their source genomes are used to create a pangenome graph. Parsimonious paths between gene families contained in the same number of genomes are used to identify co-localised gene families. This information is used to order the resulting tabular pangenome file on syntenic blocks of genes in descending order of number of genomes those blocks were present in. Gene-by-gene alignments are produced using MAFFT in order to generate a core gene alignment [14]. Installing the relevant dependencies in R allows for PIRATE to produce a pdf containing descriptive figures.

A number of supplementary tools are provided to extract, align and subset sequences, and to compare and visualize outputs. In order to facilitate integration with existing pipeline, scripts have been provided to convert the outputs of PIRATE into common formats which allows for them to be used as inputs to software used for downstream analysis, such as the PanX user-interface, SCOARY, Microreact or Phandango [6,15–17]. A full description of the methodology and comparative benchmarks has been provided in the supplementary information (Supplementary Information).

## Results and Discussion

### Benchmarking and comparison to other tools

The performance of PIRATE was assessed on a range of parameters related to its scalable application to large numbers of bacterial genomes. Three bacterial species were selected for comparison, *Campylobacter jejuni, Staphylococcus aureus* and *Escherichia coli*, representing small, medium and large pangenomes respectively (Supplementary Table 2). Memory usage and wall time were found to scale approximately linearly with increasing numbers of isolates and the amount of memory and time per sample was consistent (Supplementary Figures 1+3). PIRATE has been extensively parallelised and the availability of additional cores was found to significantly reduce runtime (Supplementary Figure 2).

**Figure 2.**
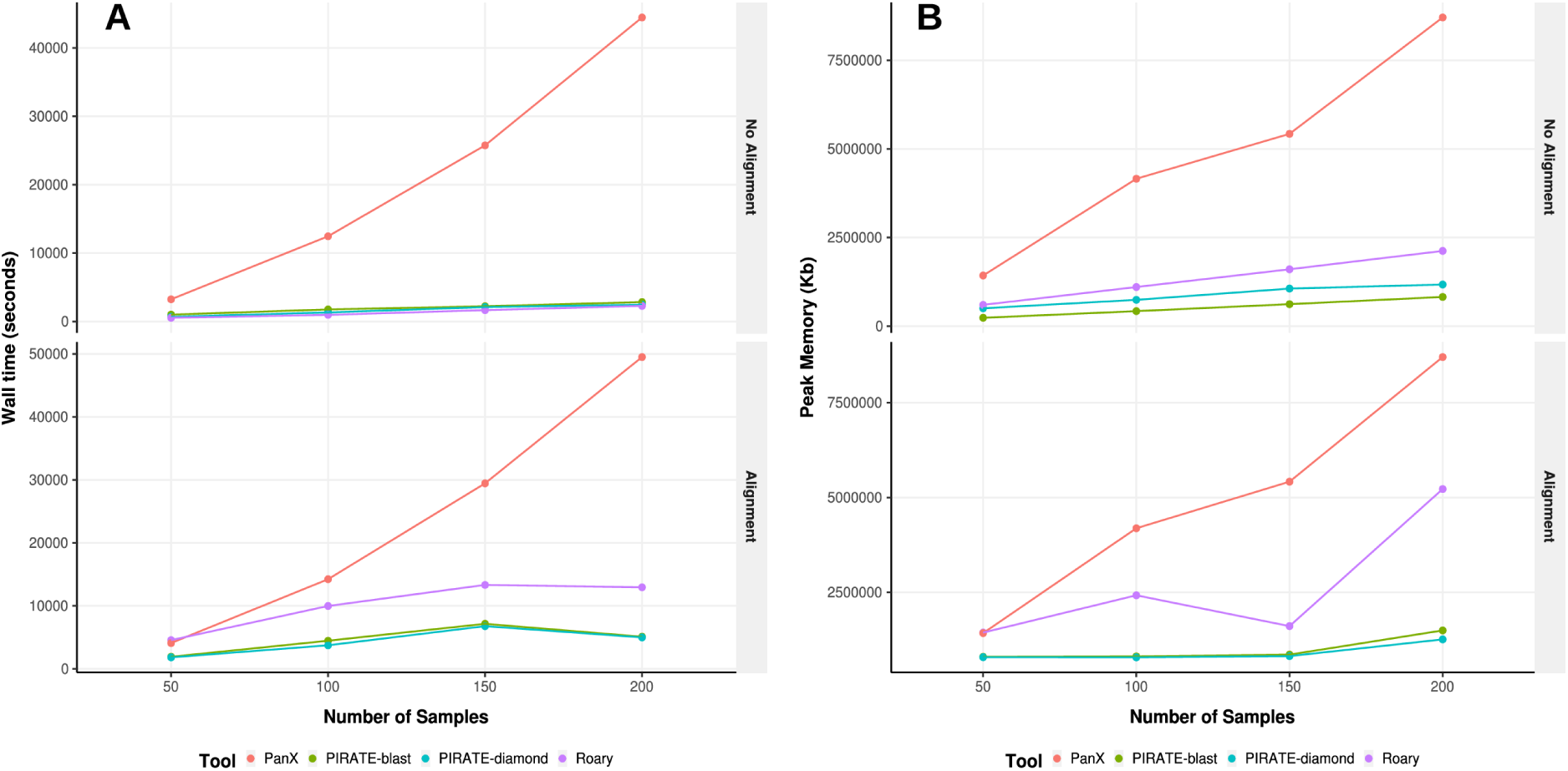
Benchmarking of PIRATE against Roary and PanX. Wall time (seconds) and peak memory usage (Kb) were recored for each tool run on a dataset of 50, 100, 150 and 200 complete *Staphylococcus aureus* genomes from the RefSeq database with and without gene-by-gene alignment

A range of tools have been developed for constructing bacterial pangenomes. For comparison, we chose two of the most widely used packages, Roary and PanX [2,6]. These tools have some similarities to PIRATE that facilitate comparison; all three tools share similar clustering workflows (BLAST/DIAMOND, MCL) and require annotated genomes as input. Differences in methodology lie primarily in the post processing of clusters, Roary uses a single percentage identity threshold for MCL clustering and separates paralogs based upon their neighboring genes and PanX splits paralogous genes using an alignment/tree-based method rather than the CDHIT-BLAST approach used by PIRATE. Each of the three tools were applied to subsets of 50, 100, 150 and 200 *Staphylococcus aureus* complete genomes downloaded from the RefSeq database (Supplementary Table 2), for comparisons on the same hardware using 8 cores [18]. It should be noted that both PIRATE and Roary include post-processing of paralogs in the comparison without alignment, producing a complete output. PanX does not do this, as alignment is a necessary step in paralog identification in this pipeline. Therefore, analyses were run with and without gene-by-gene alignment in order to make unbiased comparisons. Execution time and memory usage per sample were recorded (Figure 2).

The execution time of Roary and PIRATE scaled in an approximately linear manner with number of samples (Figure 2.A). Roary and PIRATE were faster than PanX at all time points. PanX scaled super-linearly, making application to larger datasets potentially problematic. Roary completed marginally faster than PIRATE without gene-by-gene alignment. When gene-by-gene alignment was applied both Roary and PIRATE scaled sub-linearly with number of samples. PIRATE completed substantially faster than Roary and PanX (Figure 2.A). PIRATE exhibited lower memory usage than the other tools tested, scaling sub-linearly with number of samples (Figure 2.B). In conclusion, PIRATE compared favourably in both execution time and memory usage and these metrics suggest PIRATE can be flexibly applied to large datasets on routinely available hardware.

### Application to real data

PIRATE was applied to 253 complete *Staphylococcus aureus* genomes downloaded from the RefSeq database (accessed: 08/11/18) (Supplementary Table 2) [19]. PIRATE was run on default settings over a wide range of amino acid percentage identity thresholds (45, 50, 60, 65, 70, 75, 80, 85, 90, 91-99 in increments of 1%) (Supplementary Table 2). The pangenome of *S. aureus* comprised 4250 gene families of which 2433 (57.25 %) were classified as core (>95% genomes) and 1817 (42.75 %) as accessory (Figure 3.A). Gene families with an average copy number greater than 1.25 loci per genome after paralog classification were excluded from further analysis (178 gene families, 4.18 %) as direct comparison between high copy number or potentially over-clustered families is problematic. Of the remaining 4072 gene families, 740 (18.17 %) clustered at thresholds of less than 95% percentage identity. At these thresholds a significantly different number of ‘divergent’ gene families were observed (Chi Squared test p-value = < 0.0001) between core and accessory genomes; 21.83 % of accessory genes (383/1754) clustered at less than 95% homology compared to only 15.40 % of core genes (357/2318) (Figure 3.B). A possible explanation for this is that the accessory genes may have been horizontally acquired and therefore may be from diverse genetic backgrounds with different evolutionary histories. A number of the genes exhibiting high amino acid sequence divergence have been well studied. For example, the core ‘accessory regulator’ *agr* locus exhibited a range of sequence identity clustering thresholds; *agrA* clusters at 91 %, *agrB* and *agrC* at 65 % and *agrD* at 45 % amino acid identity. This example highlights how diversification may lead to over-splitting of genes if only a single sequence identity threshold were used, even if this threshold were applicable to the vast majority of genes in the pangenome.

**Figure 3.**
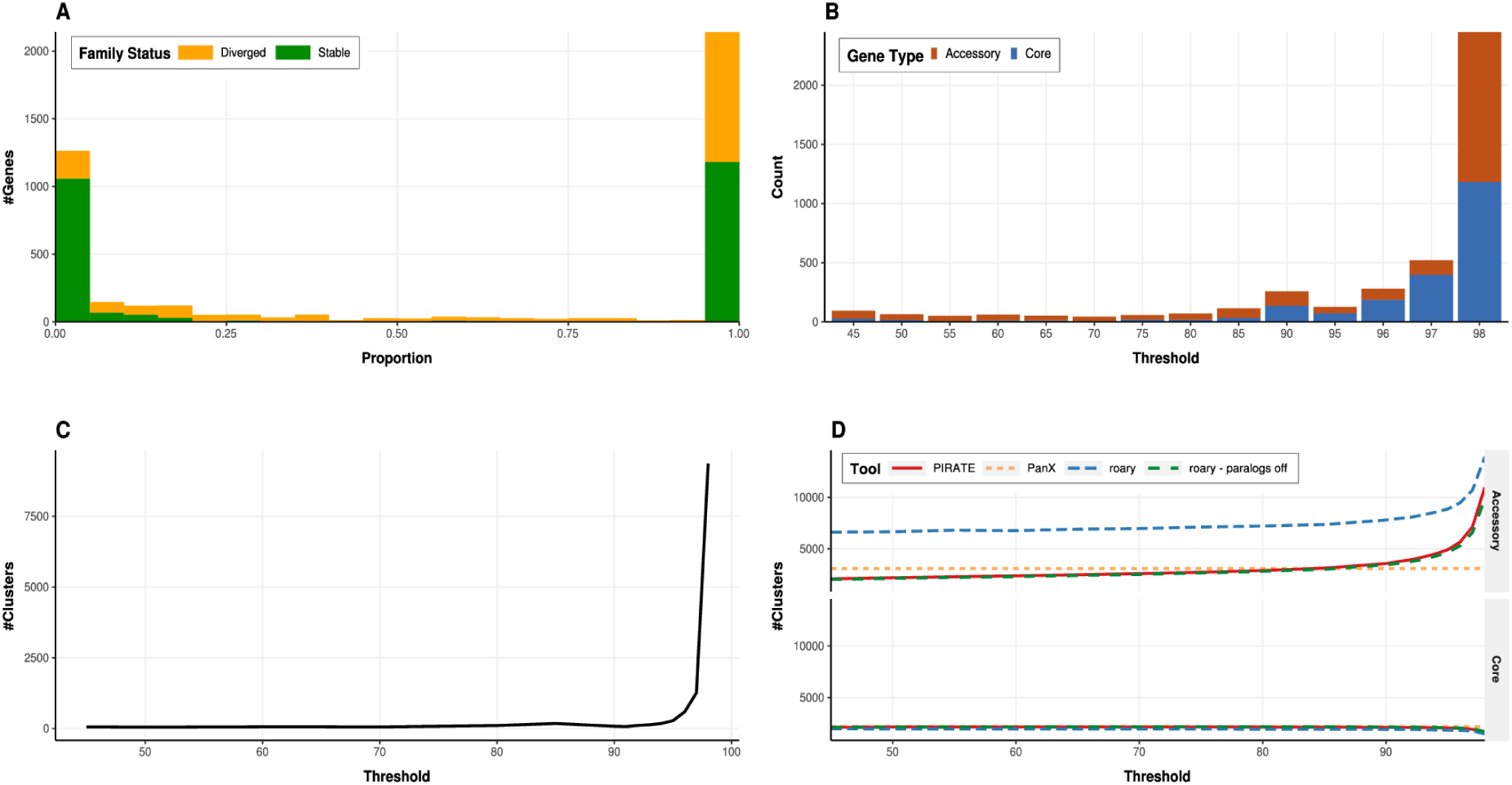
Descriptive figures of the pangenome of 253 complete Staphylococcus aureus genomes inferred using PIRATE. PIRATE was run with default parameters over a range of amino acid identity values (45-98 %). (A) The proportion of genomes in which gene families are found, indicating stable gene families (green) with a single allele at 98% amino acid identity, and diverged with >1 allele (yellow). (B) The minimum amino acid % identity cutoff at which all loci were present per gene family (core = blue, accessory = red). (C) The number of unique alleles at each amino acid percentage threshold. A unique allele is characterised as the highest percentage identity threshold at which a unique sub-cluster of isolates from a single gene family was identified by MCL. (D) Comparison of core and accessory gene/allele estimates for PIRATE (red), PanX (orange), Roary (blue) and Roary with paralog splitting switched off (green). The estimates represent ‘allelic’ variation reported by PIRATE in contrast to ‘gene content’ variation reported by the other tools. PanX provided a single estimate of core and accessory genome content as it has no analogous command to -s in PIRATE or -i in Roary to allow comparison. Core gene families are characterised as being present in greater than 95% of genomes. All tools were run on default parameters. Roary was run over a range of thresholds matching those used for PIRATE with and without paralog splitting (-s).

A steep increase in the number of unique clusters per threshold (allelic diversity) of the sample was observed at thresholds greater than 90% (Figure 3.C). At these thresholds allelic variation will begin to influence the identification of gene families in analogous tools [2,7-8]. In addition to this metric, PIRATE identifies the highest threshold at which all loci in a gene family cluster together. This value can be used to estimate the sequence similarity threshold at which an alleles is classified as a ‘gene’ by analogous tools (before paralog processing) and therefore allows for evaluation of the influence of this choice on core and accessory genome sizes (Figure 3.D). For comparison, Roary and PanX were applied to the *S. aureus* dataset (default settings). Roary was run at a range of percentage identity thresholds matching those used by PIRATE (-i option) to facilitate comparison. Paralog splitting in Roary was also switched off (-s option) to assess the influence of paralog splitting on the resulting pangenome size estimates. The number of core and accessory genes (<95% isolates) estimated by both tools was compared to those estimated using PIRATE (Figure 3.D). All tools give similar estimates of the number of core genes (PIRATE = 2141, PanX = 2191, Roary (-i 45) = 1959, Roary no paralogs (-i 45) = 2118) However, estimates of the number of accessory genes were divergent (PIRATE = 2190, PanX = 3097, Roary (-i 45) = 6620, Roary no paralogs (-i 45) = 2046). The large increase in the size of the accessory genome content inferred using Roary is primarily due to the post-processing (paralog splitting) of accessory genes. The close approximation by PIRATE of accessory content variation in Roary without paralog splitting suggests that PIRATE can be used to provide accurate estimates of pangenome composition for analogous tools before paralog splitting.

For the *S. aureus* collection the estimated number of core genes remains fairly constant at thresholds below 90% and decreases sharply at thresholds greater than 95% (Figure 3.D). This suggests that the majority of the *S. aureus* core genome would be reconstructed by tools that identify genes as clusters of sequences with >10% amino acid sequence similarity. However, the impact of more conservative thresholds on the accessory genome is pronounced. A moderate increase in the number of alleles misidentified as low frequency genes was observed at thresholds below 90% followed by a sharp increase at thresholds greater than 90%. This suggests that, even at low homology thresholds, allelic diversity in highly divergent genes inflates the number of clusters incorrectly identified as ‘accessory’ genes when using only a single homology threshold. This effect is likely to be more pronounced in organisms with large accessory genomes due to a higher number of diversified gene families in the accessory genome.

## Conclusion

Here we present PIRATE, a toolbox for pangenomic analysis of bacterial genomes, which provides a framework for exploring gene diversity by defining genes using relaxed sequence similarity thresholds. This pipeline builds upon existing tools using a novel methodology that can be applied to any annotated genomic feature. PIRATE identifies and categorizes duplicated and disrupted genes, estimates allelic diversity, scores gene divergence and contextualizes genes using a pangenome graph. We demonstrate that it compares favourably with other commonly used tools for pangenomic analysis, in both execution time and computational resources, and is fully compatible with software for downstream analysis and visualisation. Furthermore, it is scalable to multiprocessor environments and can be applied to large numbers of genomes on modest hardware. Together the enhanced core and accessory genome characterisation capability, and the practical implementation advantages, make PIRATE a potentially powerful tool in bacterial genomics - a field in which there is an urgent need for tools that are applicable to increasingly large and complex datasets.

## Acknowledgements

We would like to thank everyone who has contributed to the development of PIRATE through testing and feedback.

## Authors Contributions

S.B. developed the software and wrote the manuscript. H.A.T. and N.M.C. contributed to and tested the software. S.K.S. and E.J.F.provided guidance and contributed to the manuscript.

