## Supplementary Information for "PIRATE: A fast and scalable pangenomics toolbox for clustering diverged orthologues in bacteria"

### Performance Evaluation

The performance of PIRATE was assessed on a range of parameters related to its application to large numbers of bacterial genomes on accessible hardware. Three bacterial species were selected for comparison, *Campylobacter jejuni*, *Staphylococcus aureus* and *Escherichia coli*, representing small, medium and large pangenomes respectively.

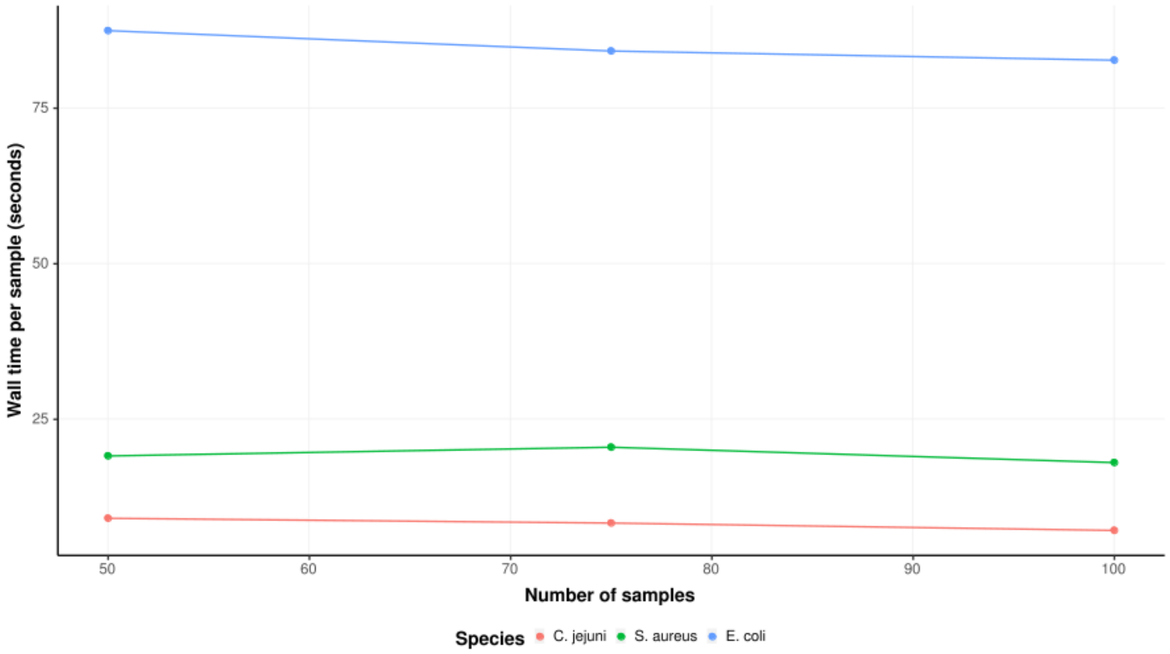


**Figure 1.** PIRATE wall time per isolate when run on comparison dataset of increasing size and complexity. The wall time per sample was calculated for subsets of 50, 75 and 100 complete genomes of *Campylobacter jejuni* (red), *Staphylococcus aureus* (green) and *Escherichia coli* (blue). PIRATE was run using default parameters.

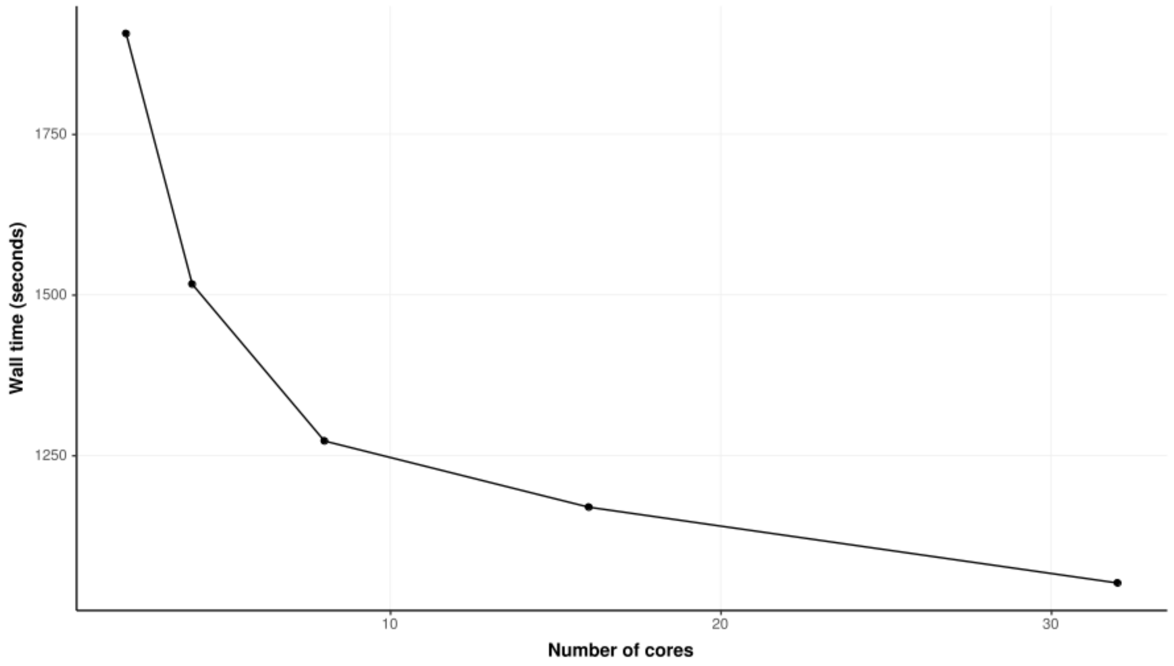


**Figure 2.** Wall time per PIRATE iteration on comparison dataset of 50 complete genomes of *S. aureus* using 2, 4, 8, 16 and 32 threads. PIRATE was run using default parameters.

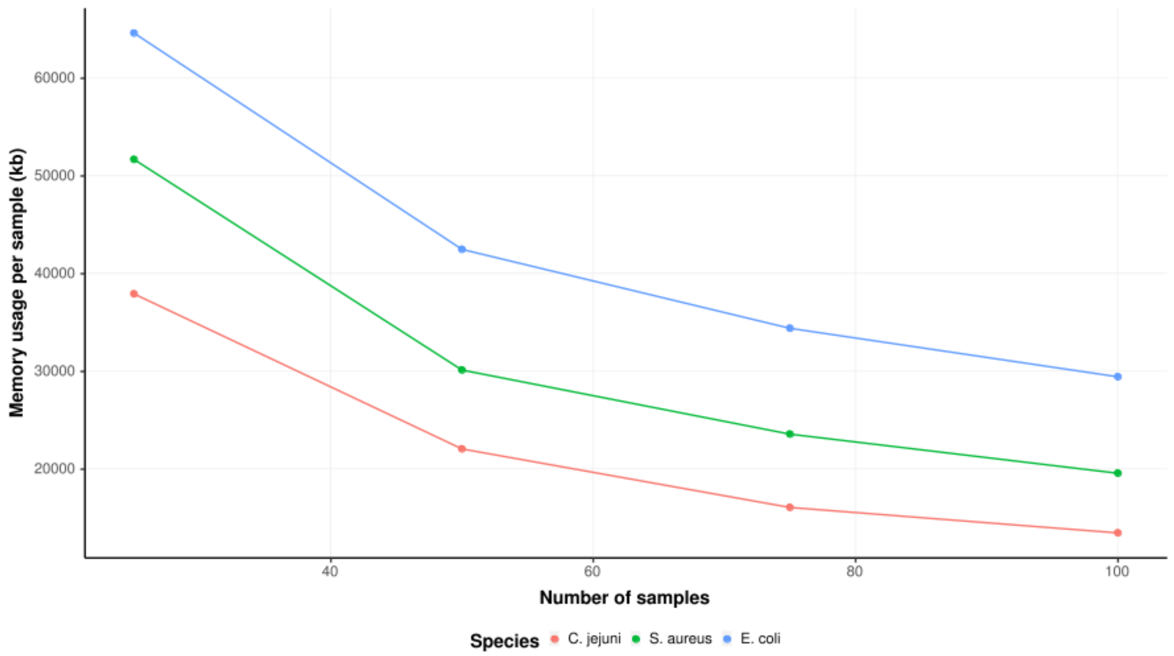


**Figure 3.** PIRATE memory usage per isolate when run on comparison dataset of increasing size and complexity. The wall time per sample was calculated for subsets of 25, 50, 75 and 100 complete genomes of *Campylobacter jejuni* (red), *Staphylococcus aureus* (green) and *Escherichia coli* (blue). PIRATE was run using default parameters for three amino acid percentage identity thresholds (50,75 and 98%).

### Method

PIRATE has been implemented in Perl and uses BioPerl for parsing files in fasta format. Parallelisation has been achieved using GNU Parallel [1]. PIRATE and all supporting scripts are available at https://github.com/SionBayliss/PIRATE under a GNU GPL 3 open-access licence. Visualisation of outputs has been achieved using the R programming language [2]. A simplified schematic of the PIRATE pipeline is presented in Figure 4.

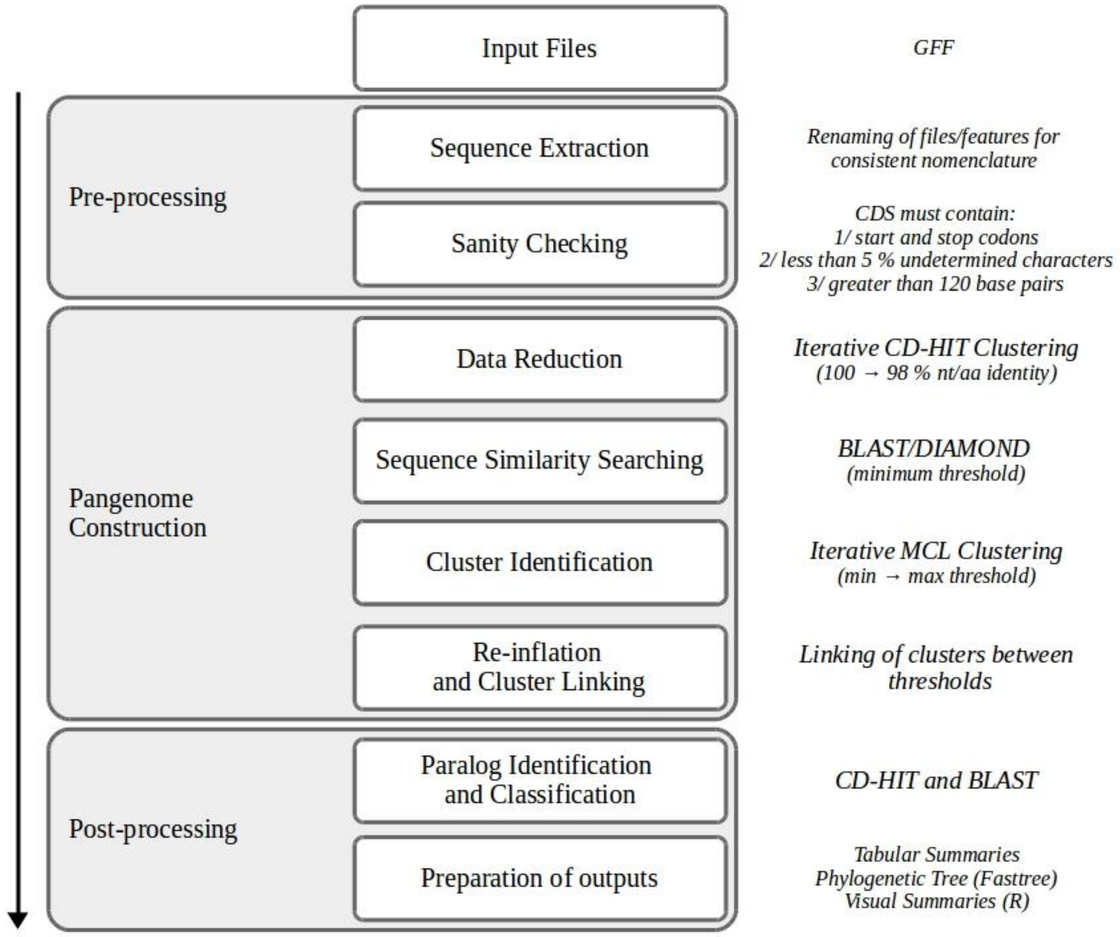


**Figure 4.** Schematic of the PIRATE pipeline.

#### *Input file format*

The input files for PIRATE are required to be in Genome Flat Format 3 (GFF3) [3]. Input files must contain nucleotide sequence information for each contig. GFF3 files produced by prokka have been extensively tested [4]. GFF3 files produced by other annotation pipelines may vary in compatibility.

#### *Input pre-processing*

Input GFF3 annotation files are parsed for the genomic features of interest, set using the --features option (CDS, tRNA, rRNA, etc.). The default features are CDS and PIRATE has been extensively tested for these features. All files with a .gff extension present in the input directory will be parsed for the feature(s) of interest. Initially, all input files are checked to ensure they adhere to the GFF3 file format and contain sequence information. Locus tag information is standardised by renaming the locus as the genome name with an ascending numeric suffix (e.g. genomeA_0001). Sequences are extracted for all features. The default behaviour is to translate nucleotide sequence for CDS features unless the --nucleotide flag is provided. Any features with greater than 5% N characters or shorter than 120 bp are excluded. CDS without start and stop codons, or with premature stop codons located within the boundaries of the gene are excluded from further analysis.

#### Sequence clustering

CD-HIT is applied as an initial clustering step in order to reduce the amount of data passed to the sequence similarity search tools [5]. CD-HIT is run on the input fasta file using an upper bounds percentage identity cutoff (option: --cdh, default: 100 %) and then an iteratively reducing this cutoff (option: --cds, default: 0.5 %) until the lower bounds cutoff value is reached or exceeded (options: --cdl, default: 98 %). Sequences which cluster at a copy number of one per genome in the collection during any of the CD-HIT iterations are considered to be ‘core’. These core CD-HIT clusters are excluded from pangenome construction and reintroduced during the cluster inflation stage. A collection of representative sequences are constructed from the longest sequences of each CD-HIT cluster. These representative sequences are passed to BLAST+ or DIAMOND for all-vs-all sequence similarity searching [6,7]. The appropriate tool is selected for the sequence type; BLASTN for nucleotide BLASTP for amino acid or, optionally, DIAMOND (--diamond). In order to facilitate multithreading the sequences are split into equally-sized chunks before the sequence similarity searching is performed on each chunk in parallel. The default parameters for sequence similarity searching is an e-value cutoff of at 1E-6 (--evalue) and dust masking is switched off. Only a single high scoring pair (HSP) is retained of each sequence comparison.

The next step applies iterative application of the Markov Cluster Algorithm (MCL) in order to cluster similar sequences at increasing stringent sequence similarity thresholds [8]. The lowest bounds percentage identity threshold (default: 50%, option: --steps) identifies initial seed clusters. HSPs with sequence similarity below this percentage identity are removed before being clustered with MCL. The resulting clusters then undergo additional rounds of MCL clustering at increasing stringent percentage identity thresholds (default: 50, 60, 70, 80, 90, 95, 98, option: --steps). Finally, the sequences previously removed by CD-HIT are reintroduced (cluster inflation) in order to generate the final clustering which include all sequences originally extracted from the input GFF3 files.

#### Gene family and allele classification

After sequence clustering, the clusters produced at the lowest similarity threshold are defined as ‘gene families’ (default: 50 %, --steps) (Figure 5). Clusterings at higher thresholds are necessarily contained within only a single gene family. These additional clustering, which represent either unique or redundant combinations of sequences, are defined as ‘alleles’ of their parent gene family. As MCL is non-deterministic, these alleles also represent possible clusters of sequences which would have been generated by an analogous pangenome tool run using a single percentage identity threshold (before paralog post-processing).

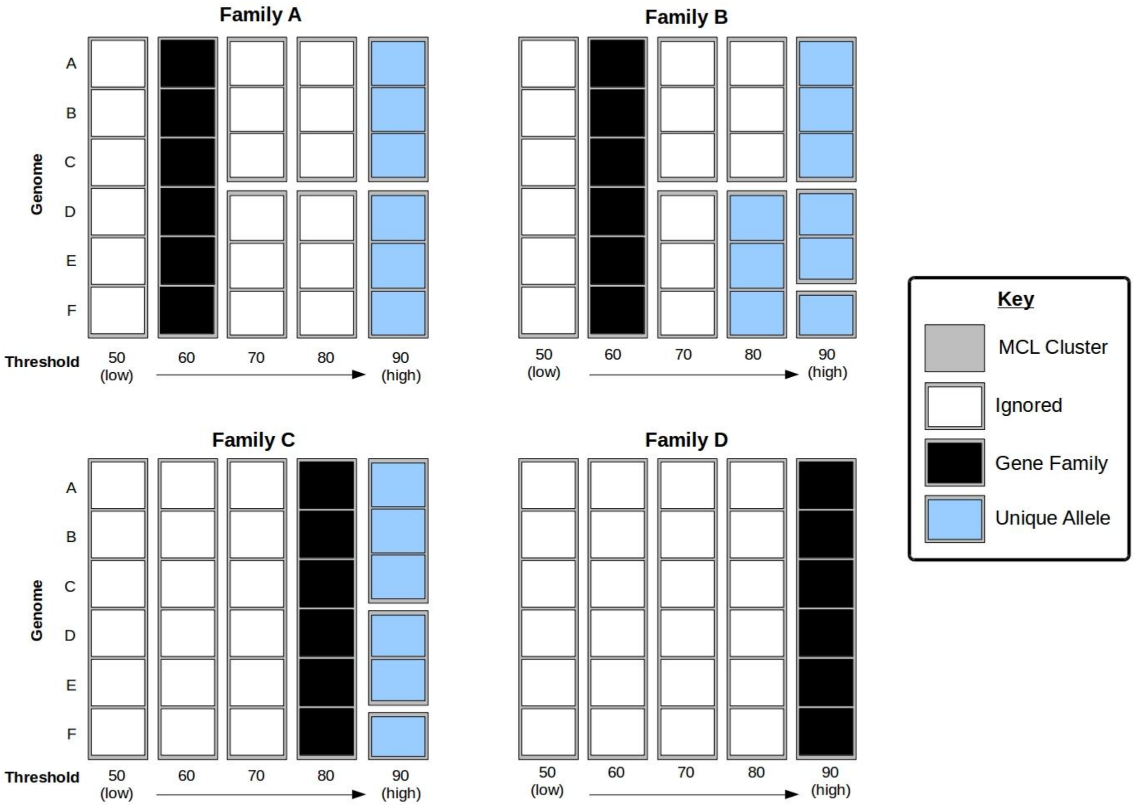


**Figure 5.** Example cluster classifications from PIRATE. White, black and blue squares represent sequences from unique genomes. Grey rectangles represent MCL clusterings of the sequences at various percentage identity cut-offs. Black squares indicate a cluster considered a 'gene family', the cluster at the highest threshold for which all sequences present in the initial clustering (lowest percentage identity) are present. Blue boxes represent 'unique alleles', clusterings at higher percentage identity thresholds which generate unique combinations of sequences from a gene family (at the higher threshold at which they are observed together). White boxes represent redundant clustering which are not output by PIRATE.

### Paralog identification and classification

Clusters which contain more than one sequence per genome undergo two additional steps - paralog classification and paralog splitting (Figure 6). Paralogous sequences are identified as either a duplication, a fission gene or both during paralog classification. Fission genes are two or more loci for which there is evidence within the dataset that they represent a single locus split into multiple loci by an indel or nonsense mutation which has caused a premature stop codon. Fission genes are identified by finding multiple non-overlapping sequences within a single genome that, when combined, have a sequence which covers the majority of another, necessarily longer, sequence within the gene family. This is performed by first identifying a series of reference clusters per gene family based on length and sequence identity by CD-HIT (>90% length, lowest threshold passed to --steps) (Figure 6.A) [5]. The longest (representative) sequence of each CD-HIT cluster is passed to BLASTN/BLASTP for sequence similarity searching [7]. Representative sequences are compared and pairs of sequences which exhibit no significant overlaps by BLAST are considered as putative fission loci (Figure 6.A). A 'scaffold' sequence is selected as the longest sequence within the gene family for which all non-overlapping loci have significant homology after filtering on the lowest threshold passed to --steps (e-value > 1E-2). All combinations of up to three non-overlapping sequences/loci are compared to the scaffold sequence in order to identify the most parsimonious coverage of the scaffold by the query sequences. This is achieved by assigning a score to the combination. The score is calculated as the number of bases of the scaffold sequence covered by the combined sequences, minus the number of bases covered more than once, minus the length any of the query sequence which does not have similarity to the scaffold sequence. Loci found to be less than two genes/features adjacent to one another on the same contig are given an adjusted score (plus 20% of scaffold length) in order to preferentially cluster adjacent loci as fission genes. Multiple sequences within a single genome exhibiting significant similarity to the scaffold and not identified as fission genes are classified as duplications.

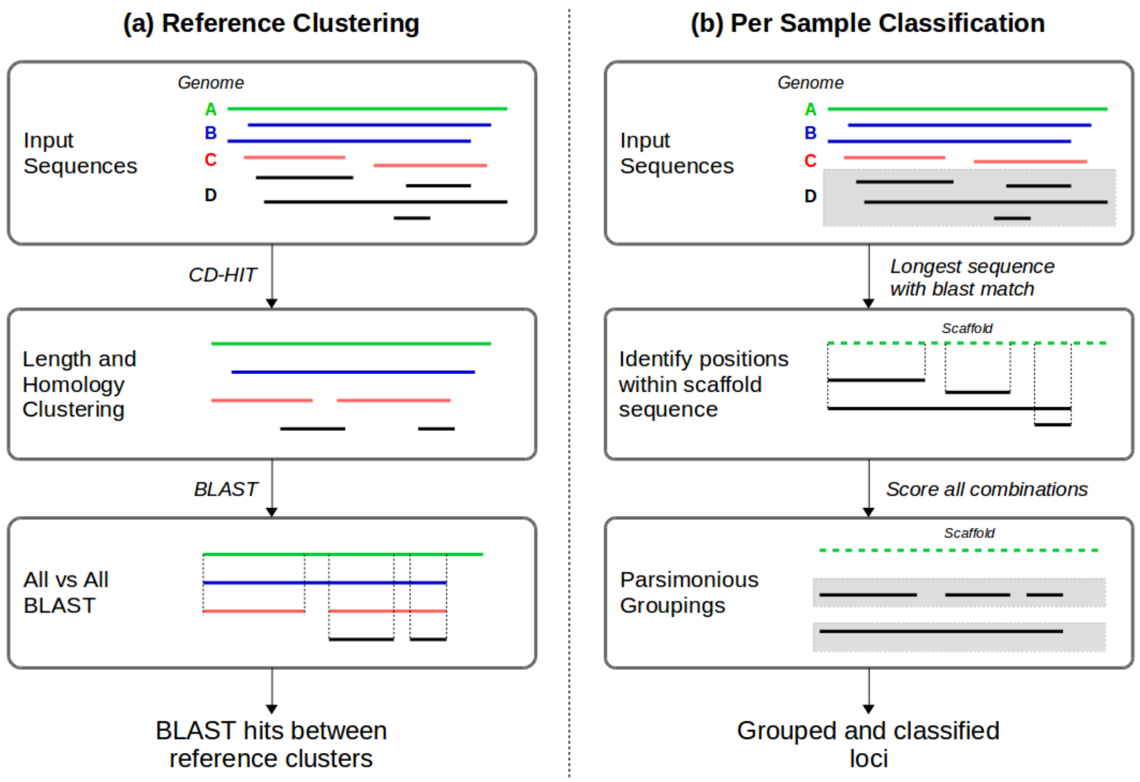


**Figure 6.** Schematic representation of the paralog classification stage of PIRATE.

### Paralog Splitting

Paralogous gene families are further analysed in order to ascertain whether there is evidence that, at any of the percentage identity thresholds, an allele might be considered *core*, *i.e.* there is one sequence per genome of the initial gene family, after considering each fission cluster as a single loci. Sequences contained within these core alleles are removed and the process repeated until no more core alleles are identified. Thereafter, core alleles are treated as separate gene families but given a shared nomenclature (gx_n: where is x is the original cluster id and n is a numeric identifier for the separated family).

### Pangenome Graph

A directed graph of the connections between the constituent loci of gene families is generated using the input GFF3 files. Paths through the graph are calculated between gene families which share only a single edge which another gene family which contains the same number of genomes. These paths represent syntenic clusters of genes and the resulting cluster order is used as an ordering variable for the gene families in the resulting output (PIRATE.gene_families.ordered.tsv). A Graphical Fragment Assembly (GFA1) file representing connection between all gene families in the pangenome is also generated and can be visualised using an appropriate visualisation suite such as Bandage [9].

### Outputs

There are a number of outputs produced by PIRATE after paralog splitting has been performed (Table 1). The STDOUT/STDERR of the PIRATE run is is logged in PIRATE.log. The gene families, after optional paralog splitting, are summarised in PIRATE.gene_families.tsv. Unique clusterings of loci are summarised in PIRATE.unique_alleles.tsv. PIRATE.pangenome_summary.txt contains a tabular count of the number of gene families present in various percentages of the input genome collection. A pseudo-fasta file is generated which denotes gene presence/absence denoted by A/T respectively in binary_presence_absence.fasta. Fast-tree is used to generate presence/absence tree in newick format (binary_presence_absence.nwk). Optionally, PIRATE will produce a series of visual summaries of the pangenome in PIRATE_plots.pdf (option: -r). A graph file is generated in GFA1 format, pangenome.gfa. The gene families file, ordered on the pangenome graph, is summarised in PIRATE.gene_families.ordered.tsv. Optionally, MAFFT alignment of each gene family can be generated using the --align option. In the case of CDS the amino acid sequence is aligned and then back-translated to nucleotide sequence, which retains the triplicate codon structure of the CDS in the resulting alignment. Aligned sequences are stored in the feature_sequences directory as both nucleotide and amino acid alignments. If the alignment option is selected PIRATE will also generate core (>95%) and full pangenome alignments with associated GFF files detailing the position of each gene in the resulting alignments (core.fasta/core.gff and pangenome.fasta/pangenome.gff). PIRATE also retains a range of files that are necessary if PIRATE needs to be re-run on the same files without recreating the pangenome clusters. Generation of these files can be disabled using the -z 0 option. Additional intermediate files can be retained by using the -z 2 option. Various scripts have been provided to convert the outputs of PIRATE into other formats for downstream analysis.

**Table 1.** Summary of output files generated by PIRATE.

| **Filename** | **Description** |
|---|---|
| PIRATE.pangenome_summary.txt | Summary statistics of pangenome |
| PIRATE.gene_families.tsv | Output annotated gene clusters |
| PIRATE.gene_families.ordered.tsv | Output annotated gene clusters ordered on the pangenome graph |
| PIRATE.unique_alleles.tsv | Unique locus clustering of gene families from PIRATE.gene_families.tsv |
| PIRATE.log | Log file |
| binary_presence_absence.fasta | Pseudo-fasta file representing the presence and absence of gene families from PIRATE.gene_families.fasta as A/T |
| binary_presence_absence.nwk | Phylogenetic tree generated from binary_presence_absence.fasta by Fasttree |
| PIRATE_plots.pdf | Summary plots (requires R and dependencies) |
| pangenome.gfa | Pangenome graph file in Genome Fragment Assembly (GFA1) format. |
| feature_sequences directory | Contains the nucleotide and, if appropriate, amino acid sequences of each gene family (--align option) |
| pangenome.fasta/core.fasta | Core (>95%) and complete pangenome alignment (--align option) |
| pangenome.gff/core.gff | Position of each gene/feature in the associated alignment (--align option). |
